# Copper deficiency alters shoot architecture and reduces fertility of both gynoecium and androecium in *Arabidopsis thaliana*

**DOI:** 10.1101/2020.06.11.146209

**Authors:** Maryam Rahmati Ishka, Olena K. Vatamaniuk

**Author notes:** Corresponding author: Olena K. Vatamaniuk.

## Abstract

Copper deficiency reduces plant growth, male fertility and seed set. The contribution of copper to female fertility and the underlying molecular aspects of copper deficiency-caused phenotypes are not well-known. We show that among copper deficiency-caused defects in *Arabidopsis thaliana* were the increased shoot branching, delayed flowering and senescence, and entirely abolished gynoecium fertility. The increased shoot branching of copper-deficient plants was rescued by the exogenous application of auxin or copper. The delayed flowering was associated with the decreased expression of the floral activator, *FT*. Copper deficiency also decreased the expression of senescence-associated genes, *WRKY53* and *SAG13*, but increased the expression of *SAG12*. The reduced fertility of copper-deficient plants stemmed from multiple factors including the abnormal stigma papillae development, the abolished gynoecium fertility, and the failure of anthers to dehisce. The latter defect was associated with reduced lignification, the upregulation of copper microRNAs and the downregulation of their targets, laccases, implicated in lignin synthesis. Copper-deficient plants accumulated ROS in pollen and had reduced cytochrome *c* oxidase activity in leaves. This study opens new avenues for the investigation into the relationship between copper homeostasis, hormone-mediated shoot architecture, gynoecium fertility and copper deficiency-derived nutritional signals leading to the delay in flowering and senescence.

**Highlight:** Copper deficiency alters shoot architecture, delays flowering and senescence, and compromises fertility by altering stigma morphology, disrupting anther lignification and dehiscence, and pollen redox status in *Arabidopsis thaliana*.

## Introduction

It has been known for decades that deficiency in the micronutrient copper in alkaline soils compromises plant fertility with the most negative impact on wheat grain production (Broadley *et al*., 2012; Graham, 1978; Graves and Sutcliffe, 1974; Shorrocks and Alloway, 1988; Solberg *et al*., 1999). In-line with the essential role of copper in plant reproduction, recent studies using synchrotron x-ray fluorescence microscopy have shown that the bulk of copper is associated with anthers and pistils in *Arabidopsis thaliana* and *Brachypodium distachyon* and failure to transport copper to reproductive structures in these species and *Oryza sativa* significantly reduces fertility (Sheng *et al*., 2019; Yan *et al*., 2017; Zhang *et al*., 2018a).

The essential role of copper stems from its participation in redox reactions and its indispensable role as a cofactor for more than 100 enzymes including those participating in important biological processes such as respiration, photosynthesis, and scavenging of reactive oxidative species (ROS) (Broadley *et al*., 2012; Burkhead *et al*., 2009; Ravet and Pilon, 2013a). It is noteworthy that because copper is redox-active, increased concentration of free copper ions causes cellular toxicity. To maintain copper homeostasis, plants regulate copper uptake by transcriptional and post-transcriptional regulation of copper transporters and *via* economizing on copper during deficiency (Burkhead *et al*., 2009; Ravet and Pilon, 2013b). The regulation of copper uptake and internal distribution include transporters from CTR/COPT and Yellow Stripe-like (YSL) families (Burkhead *et al*., 2009). Copper economy/metal switch mechanism involves the increased expression of copper-responsive microRNAs, including *miR397, miR398, miR408*, and *miR857* that, in turn, facilitate the mRNA degradation of copper-containing proteins such as Cu/Zn-superoxide dismutases (SODs), plantacyanin and laccase-like multicopper oxidases (Abdel-Ghany and Pilon, 2008; Pilon, 2017; Shahbaz and Pilon, 2019). In *A. thaliana*, copper homeostasis is controlled by a conserved transcription factor, SPL7 (Squamosa Promoter Binding Protein–like7), and a recently discovered transcription factor, CITF1 (Copper Deficiency Induced Transcription Factor 1) (Bernal *et al*., 2012; Kropat *et al*., 2005; Yamasaki *et al*., 2009; Yan *et al*., 2017). SPL7 and CITF1 function in a complex integrated pathway that is essential for copper uptake, internal transport and delivery to reproductive organs (Yan *et al*., 2017).

In addition to mineral nutrient status, other factors that determine the plant reproductive success is the timing of the transition from the vegetative to the reproductive stage and the transition to senescence. Both developmental transitions depend on diverse endogenous and environmental cues (Amasino, 2010; Cho *et al*., 2017; Johansson and Staiger, 2014; Koyama, 2018). The endogenous and exogenous cues that mediate the transition to flowering include age, hormones, photoperiod, temperature and nutrient availability, and are generally classified into five pathways: photoperiod, vernalization, age, gibberellin, and autonomous. These pathways are integrated by the florigen, Flowering Locus T (FT) that is produced in the leaf and transported *via* the phloem to the shoot apical meristem to trigger the formation of flowers (Teotia and Tang, 2015). Studies in *A. thaliana* have shown that the expression of *FT* is induced in leaves by day length/light, sucrose and its metabolite trehalose-6-phosphate (Cho *et al*., 2018; Cho *et al*., 2017; Möller-Steinbach *et al*., 2010; Srikanth and Schmid, 2011; Wahl *et al*., 2013). Apart from them, *FT* expression is also regulated by two microRNAs, *miR15(>*, and *miR172* (Cho *et al*., 2018; Cho *et al*., 2017; Möller-Steinbach *et al*., 2010; Teotia and Tang, 2015; Wahl *et al*., 2013; Wang *et al*., 2009). Our knowledge about the relationship between flowering time and copper availability is limited despite the important role of copper in plant growth and development.

Another important developmental transition that determines reproductive success is senescence. Natural leaf senescence ensures the remobilization of nutrients including minerals from senescing tissues to developing reproductive organs and seeds (Himelblau and Amasino, 2001; Leopold, 1961; Woo *et al*., 2019). Natural leaf senescence is typically triggered by the leaf age (Koyama, 2018; Woo *et al*., 2019). Environmental stresses, including nutrient deficiency, and hormones such as jasmonic acid (JA) are known to cause premature senescence (Leopold, 1961; Woo *et al*., 2019; Xie *et al*., 2014). While natural senescence increases reproductive success, premature senescence is often correlated with decreased yields. We have shown recently that JA levels increase in leaves of copper-deficient *A. thaliana* suggesting that deficiency for this mineral causes premature senescence and could be among the reasons for dropped seed yield (Yan *et al*., 2017). Hill et al., 1978 have shown however that copper deficiency delays chlorophyll degradation of mature wheat leaves and concluded that unlike other mineral deficiencies that trigger senescence, copper deficiency might, in fact, delay it. Thus, whether copper deficiency causes premature senescence or delays it, is unclear. It is noteworthy that the expression of a copper chaperone *CCH* (*ATX1-Like Copper Chaperone*) and a copper-transporting ATPase, *RAN1* (*Responsive-to-antagonist1*) is upregulated by natural senescence in *A. thaliana* pointing to the important role of copper remobilization for the reproductive success (Himelblau, 2000; Himelblau and Amasino, 2001; Himelblau *et al*., 1998).

Here, we used *A. thaliana* to perform a systematic analysis of the effect of copper deficiency on flowering and senescence that are among key developmental processes determining the reproductive success, substantiated the role of copper in male fertility and uncovered the role of copper in shoot architecture and stigma morphology.

## Materials and methods

### Plant materials and growth conditions

*Arabidopsis thaliana* (*cv*. Col-0) was used in all experiments. Plants were grown hydroponically to control copper concentrations as described in(Cho *et al*., 2017; Simpson and Dean, 2002; Yan *et al*., 2017). Copper was added at the indicated concentrations to the hydroponic solution as CuSO_4_. The standard (control) solution contained 250 nM CuSO_4_. Plants were grown in a growth chamber with the following settings: 22°C, 14-hours light/10-hours dark photoperiod at a photon flux density of 110 μmol m^−2^ s^−1^.

### Pollen germination and viability assays

Pollen grains were isolated from anthers at the stage 13-14 flower development (Sanders *et al*., 1999). For analysis of pollen grains number, 10-30 individual flowers from at least 10-30 independently-grown plants were used to manually release pollen on a media containing 0.7% (w/v) agar spread onto a microscope slide. Counting was performed after collecting images on the Axio Imager M2 microscope (Zeiss).

Pollen viability was evaluated using fluorescein diacetate (FDA) according to (Bou Daher *et al*., 2009). Briefly, a 10 mg/mL FDA stock was prepared in acetone and stored at −20°C. A working solution of FDA was made by diluting FDA stock in 10% sucrose solution to a final concentration of 0.2 mg/mL. Pollen grains were released from five to ten open flowers by tapping anthers into 200 μl FDA solution followed by 5 minutes incubation in the dark. An aliquot (50 μl) was then transferred to a microscope slide and viable pollen was analyzed by fluorescence microscopy using the Axio Imager M2 microscope (Zeiss) and the FITC filter set.

Pollen germination was analyzed as described in (Fan *et al*., 2001). Briefly, pollen grains were spread on pollen germination media for overnight growth at 25°C. After 24 hours incubation, the germinated pollen grains were imaged using the Axio Imager M2 microscope (Zeiss) and counted using ImageJ software. For the copper rescue, CuSO_4_ was added directly to the cooled pollen germination media (50°C) to a final concentration of 20 nM. For the L-ascorbate rescue experiment, L-ascorbate was added directly to the cooled pollen germination media (50°C) to a final concentration of 5 μM.

### Analysis of anther dehiscence

Anther dehiscence was analyzed according to (Yan *et al*., 2017). In brief, anthers from stage 14 flower development were analyzed using a Leica S6E stereomicroscope at 40X magnification. For plants grown without CuSO_4_, 50 flowers from ten independently-grown plants were analyzed. For control condition, 20 flowers from ten independently-grown plants were analyzed.

### Preparation of ultra-thin flower cross sections

The effect of copper deficiency on anther morphogenesis was analysed using light microscopy on semi-thin sections using the procedure modified from (Zhao *et al*., 2002). Briefly, the entire primary inflorescences containing floral buds and flowers at different developmental stages were fixed for three hours in 2% glutaraldehyde (v/v) in 0.05 M cacodylate buffer (pH 7.4). After washing three times, 10 minutes each time, in cacodylate buffer, samples were dehydrated for 10 min in ethanol series of 25%, 50%, 70%, 85%, and two times in 100%. The molecular sieve was used in 100% ethanol to trap any extra water that may have been accumulated in ethanol. Samples were then placed in 100% absolute acetone with molecular sieve for two more changes, 10 minutes each. All the above-mentioned steps were conducted on ice. Inflorescences were dissected into individual buds and flowers prior to transferring to epoxy resin (Quetol 651, catalogue # 14640 VWR) series. Individual buds/flowers were then embedded in epoxy : acetone (1:3 ratio) for four to eight hours, transferred to epoxy: acetone (1:1 ratio) for four to eight hours, transferred to epoxy: acetone (3:1 ration) for eight hours, and 100% epoxy for 12 hours (or overnight). Finally, dissected samples were polymerized in molds at 60°C for 12 hours. Ultra-thin cross sections (1 μm) were obtained using an ultramicrotome (Leica-Ultracut-UCT) equipped with a diamond knife. Sections were then heat fixed on glass slides for about 15 minutes. Obtained sections were stained with 0.5% toluidine blue (Sigma-Aldrich) for one minute and rinsed three times with deionized water. Images were taken using Axio Imager M2 microscope (Zeiss).

### Lignin staining

Anthers were dissected from flowers at stage 13 to 14 of flower development according to (Sanders *et al*., 1999) and stained with phloroglucinol-HCl (Hao *et al*., 2014b). The staining solution was prepared by mixing two parts of 2% (w/v) phloroglucinol in 95% (v/v) ethanol with one part of concentrated HCl. For visualizing lignin in the inflorescence stem, transverse hand sections of stems were fixed in a solution comprised of three parts absolute ethanol to one part acetic acid for 15 minutes, then rinsed with deionized water and stained with phloroglucinol - HCl. Samples were imaged 10 minutes after staining using the Axio Imager M2 microscope (Zeiss, Inc). Images were collected with the high-resolution AxioCam MR Camera and processed using the Adobe Photoshop software package, version 12.0.

### ROS staining of pollen grains using CM-H2DCFDA

ROS staining was conducted as described in (Muhlemann *et al*., 2018). In brief, mature pollen grains from plants grown hydroponically with or without 250 nM CuSO_4_ were released into the pollen germination media (as defined in Fan, Wang et al. 2001) and 5 μM CM-H2DCFDA (Thermo Fisher). Pollen grains were then incubated at 28°C for 20 minutes and pelleted by a quick spin. The staining solution was then replaced with pollen germination media followed by imaging using FITC filter set on the Axio Imager M2 microscope. In order to quantify the signal intensity, the background signal was determined for each sample using unstained pollen grains. Pollen grains showing signal intensity above the background level were then counted.

### Cytochrome *c* oxidase enzyme activity

Plants were grown hydroponically with or without 250 nM CuSO_4_ for four weeks. Mitochondria were isolated from rosette leaves using the procedure described in (Keech *et al*., 2005). Cytochrome *c* oxidase activity was measured using Cytochrome *c* Oxidase Assay Kit (Sigma-Aldrich, catalog number CYTOCOX1) according to the manufacturer’s instruction. This assay was done using two independent experiments.

### Isolation of total RNA and RT-qPCR

Tissues were collected from plants grown hydroponically at the indicated copper concentrations, flash-frozen in liquid nitrogen, and stored at −80°C prior analyses. All samples were harvested between seven and eight Zeitgeber time, unless otherwise stated. Total RNA was isolated using TRIzol reagent (Invitrogen) according to the manufacturer’s instructions. One microgram of total RNA was then treated with DNase I (New England Biolabs) prior to the first-strand cDNA synthesis using AffinityScript RT-qPCR cDNA synthesis kit (Agilent Technologies). RT-qPCR analysis was conducted using iQ SYBRGreen Supermix (Bio-Rad) according to manufacturer’s instructions in the CFX96 real-time PCR system (Bio-Rad). *AtACT2* (AT3g18780) was used as a reference gene for data normalization. RT-qPCR experiments were conducted using three independent experiments, each with three technical replicates. The list of oligos is shown in **Supplementary Table S1**.

## Results

### Copper deficiency increases shoot branching that can be rescued by auxin

We first tested the effect of different copper concentrations on the growth and development of *A. thaliana* because this has not been done comprehensively. We grew plants hydroponically in different CuSO_4_ concentrations ranging from zero (no copper added) to 500 nM. As would be expected due to the essential nature of copper, plants grown under low copper were smaller in stature (**Supplementary Fig. S1A**) and had a smaller size of rosette leaves (**Supplementary Fig. S1B**) compared to plants grown under control conditions (i.e., 250 nM CuSO_4_). Unexpectedly, plants grown under copper deficiency (i.e., 0 to 10 nM) developed more axillary branches (**Fig. 1A**) and had a longer primary inflorescence upon transition to flowering (**Fig. 1B**) compared to plants grown under control condition. We also noted that apical flower bud on the primary inflorescence was aborted in plants grown under copper deficiency (**Fig. 1C-I, II**), suggesting that increased branching might be related to the removal of auxin-dependent apical dominance.

**Fig. 1.**
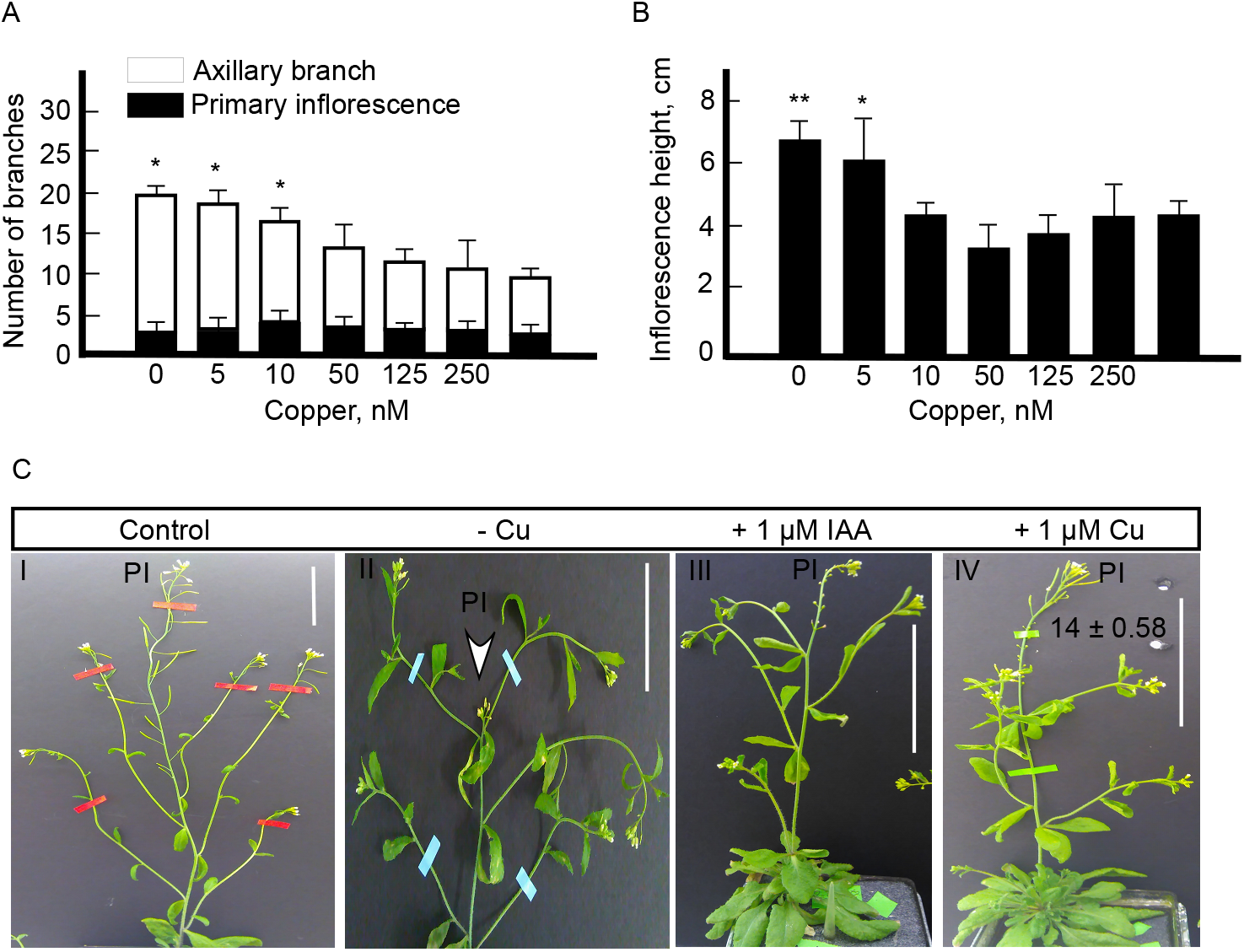
Copper deficiency alters shoot architecture in Arabidopsis. (A) Plants were grown hydroponically under indicated copper concentrations. The number of primary inflorescences and axillary branches was evaluated at late reproductive stage (stage 8 of *A. thaliana* development as determined by (Clark, 2001)). (B) shows the height of primary inflorescence upon transition to flowering (upon opening of the first flower on primary inflorescence) in plants grown under different copper concentrations. In (A) and (B) values are mean ± SE (n = 3 independent experiments with 5-10 plants analyzed in each experiment). Asterisks (* and **) indicate statistically significant difference compared to control condition (i.e., 250 nM Cu) with *P* < 0.05 and < 0.001, respectively, using Student’s *t*-test. (C) Shows representative images of plants grown hydroponically with or without 250 nM CuSO_4_. Panel C-I, control, shows a representative image of a plant grown hydroponically with 250 nM copper. Panel C-II, - Cu, shows a representative image of a plant grown hydroponically without copper supplementation. Note a partially aborted apical flower, while increased branching occurred at late reproductive stage. Panel C-III, + 1 μM IAA, shows a plant grown hydroponically without CuSO_4_ for three weeks followed by application of 1 μM IAA (auxin) onto rosette leaves and apical meristem for five days. Panel C-IV, + 1 μM Cu, shows a representative image of a plant grown hydroponically without CuSO_4_ for three weeks followed by the application of 1 μM CuSO_4_ onto rosette leaves and apical meristem for five days. Deionized water was used for mock treatment in both auxin and copper rescue experiment. Both copper and auxin rescue experiments were done at least three independent times, with three to five individually grown plants used each time. Scale bar = 1 cm in all.

Consistent with this suggestion, exogenous application of indole-3-acetic acid (IAA) to the shoot apex has decreased shoot branching (**Fig. 1C-III**). Exogenous application of copper to the shoot apex also partially rescued shoot branching of copper-deficient plants (**Fig. 1C-IV**). These data link copper to auxin signaling in establishing inflorescence architecture. It is noteworthy that exogenous copper but not IAA has partially rescued seed set of copper-deficient plants, signifying the role of copper in reproduction. Plants that were grown at 500 nM CuSO_4_ did not show any distinguishable phenotypic deviation from those that were grown under 50, 125 and 250 nM CuSO_4_. The concentration of 250 nM was used as a control for comparisons from here on.

### Copper deficiency delays transition to flowering, reduces the number of flowers and alters the expression of *FT* and *miR172*

It has been shown previously that *Chrysanthemum morifolium* grown under copper deficiency flowers later compared to copper-sufficient plants (Graves and Sutcliffe, 1974). We noticed that copper-deficient *A. thaliana* flowers later too. Here, we tested whether the late flowering time is caused by the delayed vegetative-to-reproductive stage transition or the slower growth rates of plants under copper deficiency. The time to flowering and the rosette leaf number upon transition to flowering are used as common indicators for monitoring the time from the vegetative-to-reproductive stage transition in *A. thaliana* (Pouteau and Albertini, 2009). We found that copper deficiency not only delayed the time to flowering but also increased the number of rosette leaves upon transition to flowering and thus, delayed the transition from the vegetative to the reproductive stage (**Fig. 2A-B**). By contrast, the number of floral buds on the primary inflorescence upon transition to flowering was significantly reduced in copper-deficient plants (**Fig. 2C**).

**Fig. 2.**
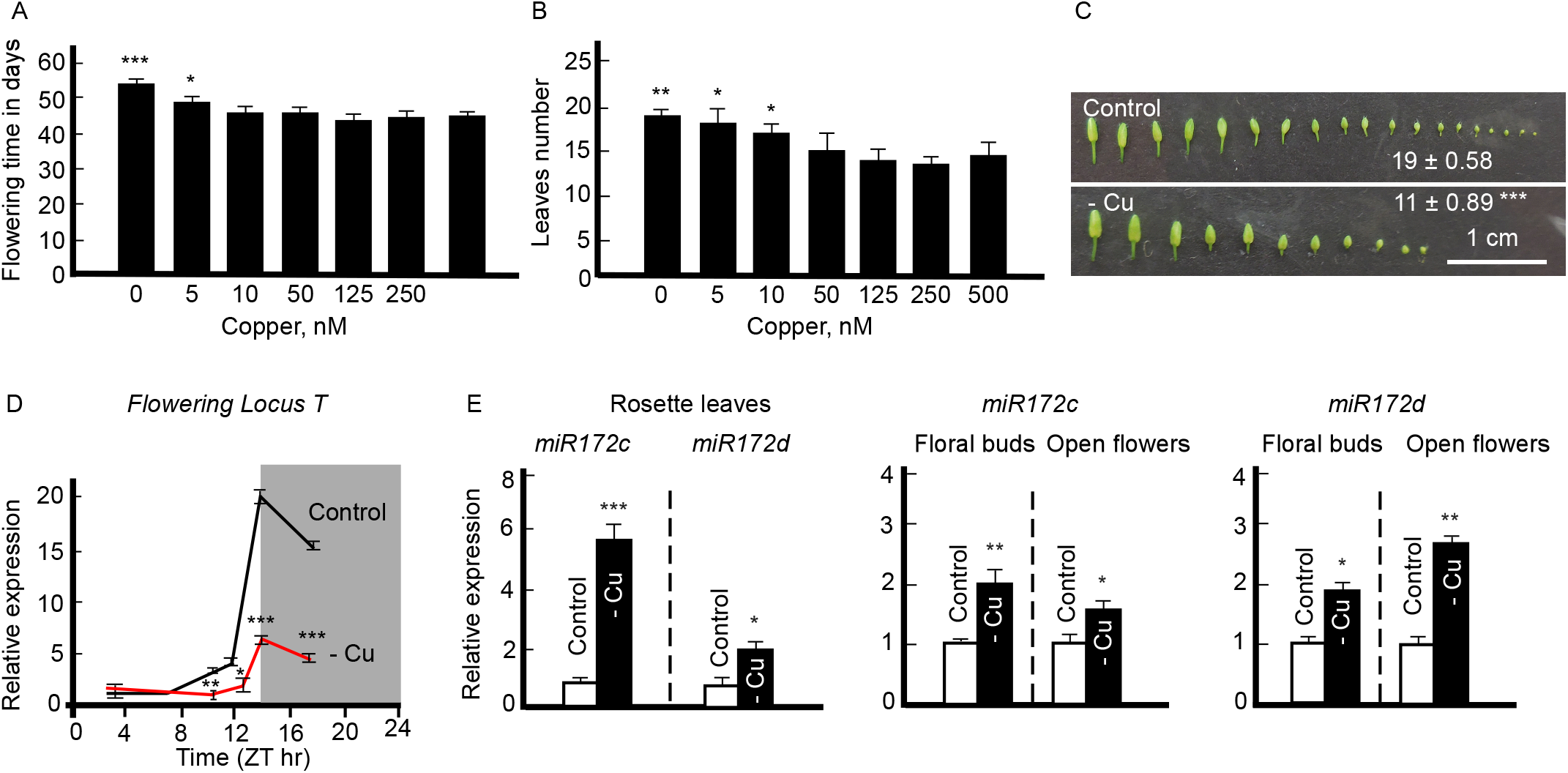
Copper deficiency delays transition to flowering by altering the expression of *FT* and *miR172* in Arabidopsis. Flowering time (A), number of rosette leaves (B), and number of floral buds (C) are shown upon transition to flowering (A and B) or 5 days after transition to flowering in (C) for plants grown hydroponically under indicated CuSO_4_ concentrations. In (C), plants were grown hydroponically with or without 250 nM CuSO_4_. (A) to (C) Values are means ± SE (n = 3 independent experiments with 5-10 plants analyzed in each experiment). Asterisks (*, **, and ***) indicate statistically significant differences vs. controls with *P* < 0.05, 0.001, and 0.0001, respectively, using the Student’s t-test. (D) Diurnal time course expression of *Flowering Locus T* (*FT*) in four-week-old rosette leaf. Plants were grown hydroponically with or without 250 nM CuSO_4_ for four weeks. Samples were collected based on Zeitgeber time, where the Zeitgeber hour one is the first hour of light after the dark period. (E) Transcript abundances of *miR172c* and *d* in four-week-old rosette leaf and flower samples. For analysis using rosette leaves, plants were grown hydroponically with 250 nM CuSO_4_ for four weeks and then transferred to a fresh medium lacking copper and grown for one week to introduce copper deficiency. (D) to (E) show mean values ± SE (n = 3 independent experiments with at least three to five plants analyzed in each experiment). Asterisks, * (*P* < 0.05), ** (*P* < 0.01) and *** (*P* < 0.001), indicate statistically significant differences compared to control condition using the Student’s t-test.

The transition to flowering is associated with the expression in the rosette leaves and transport to the shoot apical meristem of the floral activator, *Flowering locus T* (*FT*) and a floral identity marker, *miR172* (Duan, Wei *et al*., 2017; Hofmann 2017; Díaz□ Manzano, Cabrera *et al*., 2018). We, thus, hypothesized that the delayed vegetative-to-reproductive stage transition in *A. thaliana* under copper deficiency can be caused by the decreased expression of *FT* and *miR172*. As we predicted, the transcript abundance of *FT* was significantly reduced at ZT10 (Zeitgeber time 10), ZT12, and ZT14 in leaves of plants grown without added copper compared to plants grown under control conditions. This reduction continued even through the dark cycle (**Fig. 2D**).

We then tested *miR172* expression in both rosette leaves and flowers (**Fig. 2E**). Among the five isoforms tested, we could only detect isoforms *c* and *d*. We found that *miR172c* and *d* transcript abundances were significantly increased in both rosette leaves and flowers of copper-deficient plants (**Fig. 2E**). Together, these results suggest copper-deficiency-driven delayed flowering is, in part, mediated by changes in the expression of *FT* and is independent of *miR172*.

### Copper deficiency reduces fertility and impacts both reproductive organs

Our analysis of copper localization in the reproductive organs of *A. thaliana* has shown that copper is associated with both anthers and pistils (Yan *et al*., 2017). This finding suggested copper might be needed for the fertility of both reproductive organs. To test this prediction, fertility and seed production were evaluated using reciprocally crossed wild-type *A. thaliana* grown with or without copper. At the onset of this study, we evaluated the effect of different concentrations of copper on seed production. As would be expected, plants were sterile when grown without copper supplementation (**Fig. 3A- B**). Increasing copper concentration in hydroponic medium improved fertility (**Fig. 3B**).

**Fig. 3.**
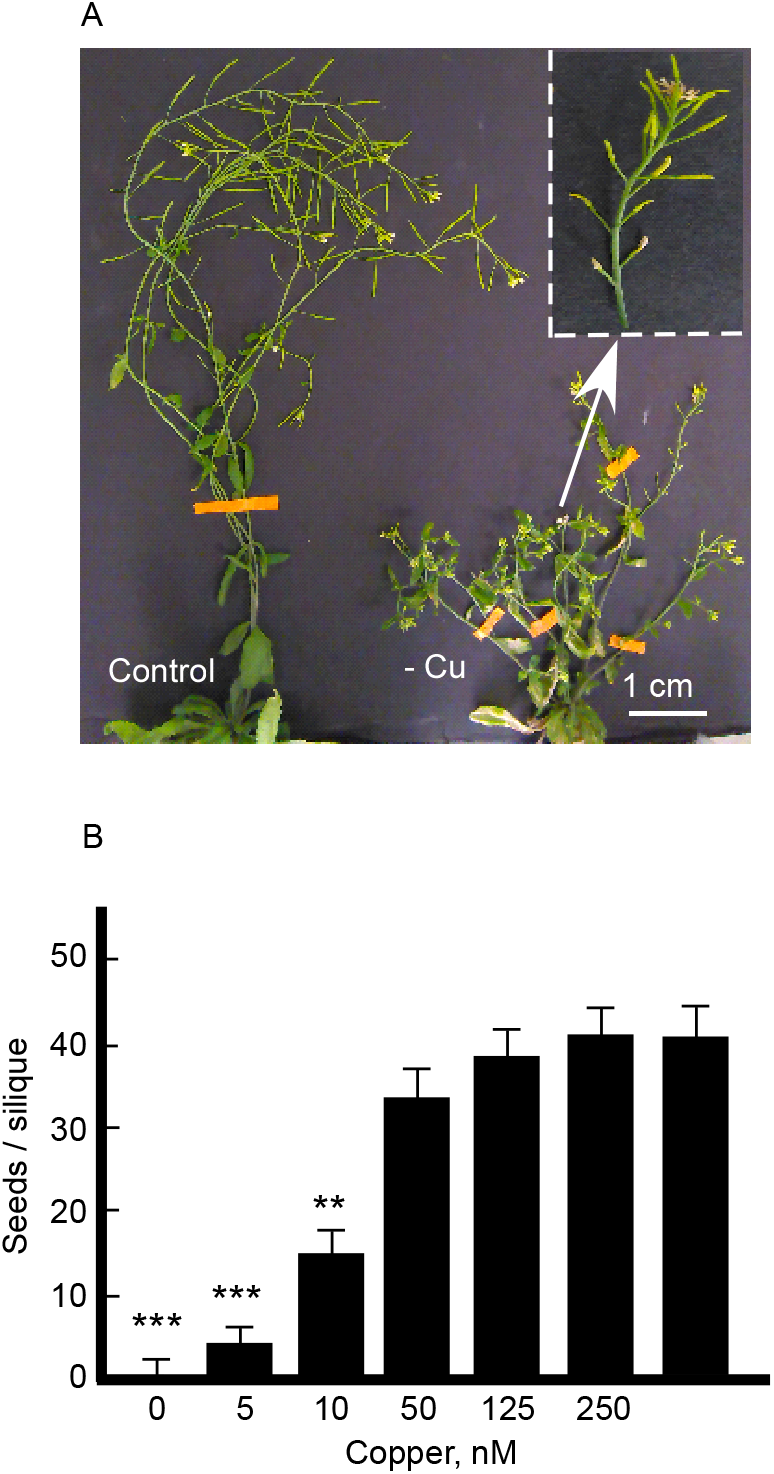
Copper deficiency reduces fertility in both reproductive organs. (A) Shows representative images of plants grown hydroponically with or without 250 nM CuSO_4_ until reproductive stage. Inset shows magnified primary inflorescence of a copper-deficient plant which is infertile. (B) Seeds number per silique are shown for plants grown under different copper concentrations as indicated. Values are mean ± SE (n = 3 independent experiments with 5-10 plants analyzed in each experiment). Asterisks (** and ***) indicate statistically significance difference compared to control condition (i.e., 250 nM Cu) with *P* < 0.001 and 0.0001, respectively, using the Student’s t-test.

To find which reproductive organs, male or female, or both were affected by copper scarcity, reciprocal crosses were conducted between plants grown under control condition (250 nM CuSO_4_) *vs*. plants grown without copper supplementation. As shown in **Table 1**, the seed production was severely affected when gynoecium of plants grown under control conditions was fertilized with pollen from plants grown without copper. However, when the gynoecium of a copper-deficient plant was a recipient of pollen from a plant grown under the control condition, seed production was completely abolished (**Table 1**). Together, these results indicated that copper deficiency causes defects in both reproductive organs with the most pronounced defect attributed to the gynoecium.

**Table 1.** Copper deficiency reduces fertility of both androecium and gynoecium. Reciprocal crosses were conducted between *A. thaliana* plants grown with or without 250 nM CuSO_4_. Statistical significant levels were determined by the Pearson’s Chi-squared test (*x*^2^) with a minimum threshold set at *P* < 0.05 compared to reciprocally-crossed plants grown under control conditions.

| Cross | Average of seed<br>#/silique ± SE | Total # of siliques<br>analyzed | *P*-value |
| --- | --- | --- | --- |
| Female control x male control | 24 ± 2.02 | 3 | 1 |
| Female control x male 0 Cu | 11 ± 3.13 | 8 | <0.01 |
| Female 0 Cu x male control | 0 | 22 | <0.01 |

### Copper deficiency reduces stigmatic papillae formation

To evaluate the role of copper in the gynoecium fertility, we took a closer look at pistils and noticed that the stigma in almost 90% of the copper-deficient plants lacked or had shorter papillae (**Fig. 4A**). Because stigmatic papillae serve as attachment sites for pollen and are required for fertilization (Kang *et al*., 2003; Thorsness *et al*., 1993), we speculate that papillae length reduction or complete abolishment under copper deficiency is a major contributing factor to the defect in female fertility (**Table 1**).

**Fig. 4.**
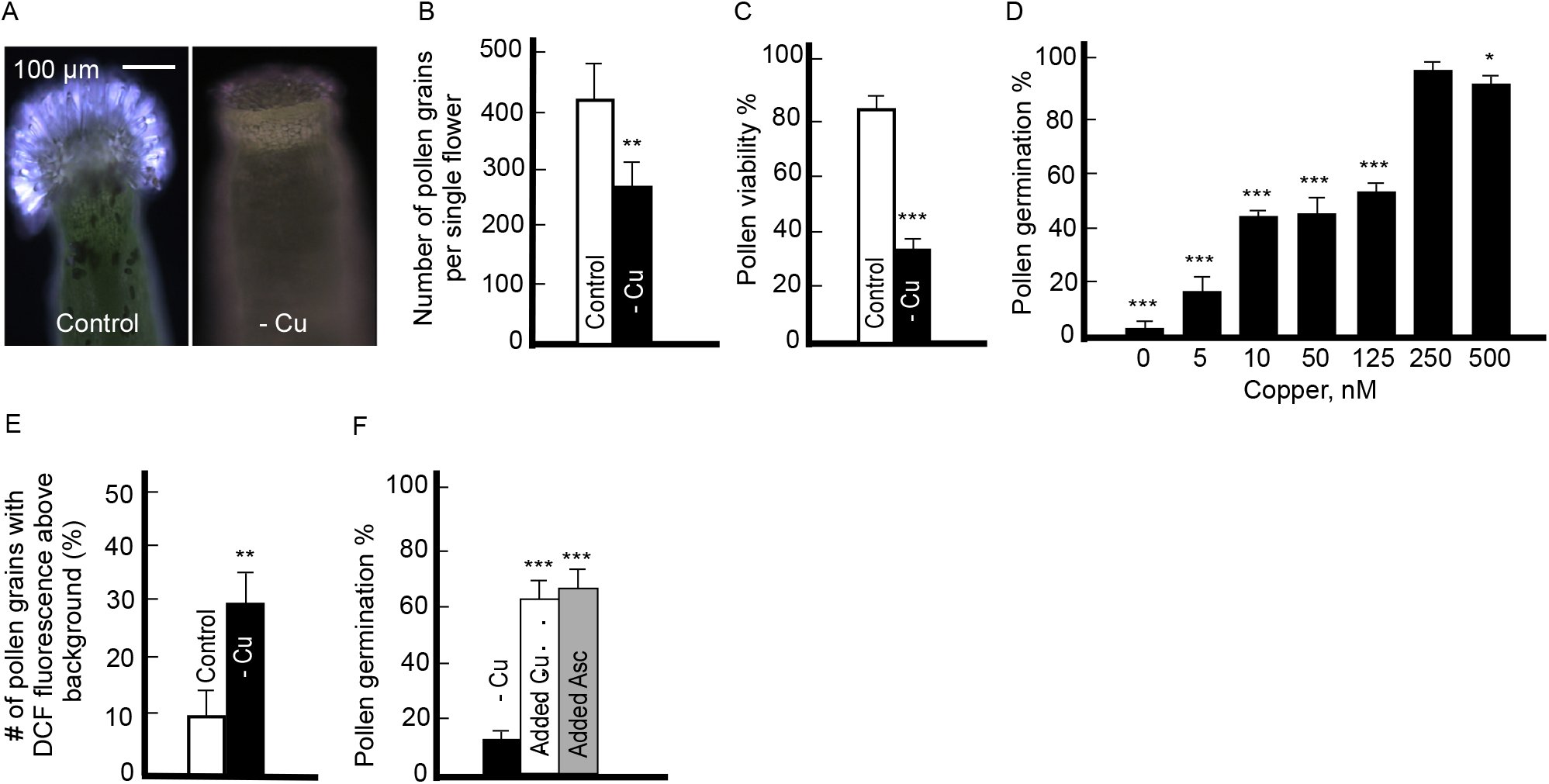
Copper deficiency dramatically reduces stigma and pollen grains fitness. (A) Shows representative images of the light microscopy of the stigma from plants grown hydroponically with or without 250 nM CuSO_4_. Note that stigma from a copper-deficient plant is almost papilla-less. (B) Number of pollen grains per single flower is shown. (C) to (D) Pollen viability and *in vitro* germination. (E) ROS level in pollen grains. Graph represents DCF fluorescence quantification in pollen. (F) Application of 20 nM copper or 5 μM L-ascorbate directly to the pollen germination media rescues defect in pollen germination of Cu-deficient plants. In (B) to (E), plants were grown hydroponically in different concentrations of CuSO_4_ as indicated. (B) to (F) Values are mean ± SE (n = 3 independent experiments with 5-10 plants analyzed in each experiment). Asterisks (*, **, and ***) indicate statistically significant differences compared to control condition in (B) to (E) or copper-deficient condition in (F) with *P* < 0.05, 0.01 and 0.001, respectively, using the Student’s t-test.

### Copper deficiency reduces pollen number, viability, and germination while increasing ROS production

Concerning the reduced male fertility (**Table 1**), copper-deficient plants produced fewer pollen grains (**Fig. 4B**), more than 60% of which was unviable compared to plants grown under control condition (**Fig. 4C**). Decreasing copper concentration in the plant growth medium also decreased pollen germination and nearly abolished it when plants were grown without copper (**Fig. 4D**). Because copper deficiency causes oxidative stress, we asked whether observed pollen defects could be associated with the accumulation of reactive oxygen species (ROS). To evaluate ROS level in pollen grains, we used the general ROS sensor, 5-(and 6)-chloromethyl-2’,7’-dichlorodihydrofluorescein diacetate (CM-H2DCFDA), which is converted to the highly fluorescent 2’,7’-dichlorofluorescein (DCF) upon oxidation by cellular ROS (Muhlemann *et al*., 2018). We found that pollen from copper-deficient plants accumulated more ROS compared to plants grown under control conditions (**Fig. 4E**). The exogenous application of an antioxidant, L-ascorbate, or copper directly to the pollen germination medium increased pollen germination by almost six folds (**Fig. 4F**). Together, these results suggest that at least some of the pollen fertility defects of copper-deficient plants, namely, pollen germination, are due to copper deficiency-promoted oxidative stress.

To identify other copper-dependent contributors to pollen fertility, we examined the activity of cytochrome *c* oxidase (COX), a copper-containing protein complex that plays a crucial role in the mitochondrial respiratory chain, and is essential for the cellular energy production (Droppa *et al*., 1984). Due to difficulty in collecting enough pollen material for the isolation of mitochondria, we measured COX activity in mitochondria that were isolated from rosette leaves of plants grown with or without 250 nM copper. As would be expected because of the essential role of copper in COX function, copper deficiency decreased COX activity by almost three-fold compared to that in the control condition (**Table 2**). These results suggested that the observed fertility defects might be, in part, due to the decreased cellular energy production.

**Table 2.** Copper deficiency reduces cytochrome *c* oxidase activity in rosette leaves of *A. thaliana*. Plants were germinated and grown hydroponically with or without 250 nM CuSO_4_ for four weeks. Mitochondria were extracted according to Keech et al. (2005). This experiment was done twice, each with three biological replicates, with both experiments showing similar results. Representative of two experiment was shown here as mean values ± SD. *** indicates statistical significance difference compared to control condition with P < 0.0001, using the Student’s *t*-test.

| Conditions | Cytochrome c oxidase (units/ml) |
| --- | --- |
| 250 nM copper | 1151 ± 30.5 |
| 0 nM copper | 361 ± 10.5 *** |

### Copper deficiency compromises anther and stigma cell specification in *Arabidopsis*

To gain further insight into the effect of copper deficiency on reproductive organs, Ultra-thin transverse sections were prepared from the *A. thaliana* floral buds collected at different developmental stages from plants grown with or without 250 nM CuSO_4_. The anther wall of plants grown under copper sufficient condition contained four defined cell layers including epidermis, endothecium, middle layer, and tapetum (**Fig. 5A-I, -II and -III**). However, the wall of anther lobes in copper-deficient floral buds looked contorted without any defined cell layers in all developmental stages, as early as the stage of pollen mother cell formation (**Fig. 5A-V**). The same undefined structures remained throughout the middle and late stages of anther development in copper-deficient plants (**Fig. 5A-VI, -VII**). Similarly, a cross-section through gynoecium of the copper-deficient flowers did not show defined structures (**Fig. 5A-VIII**) compared to that in the control condition (**Fig. 5A-IV**). Together, these observations indicate that copper deficiency adversely affects the morphology of both androecium and gynoecium in their development.

**Fig. 5.**
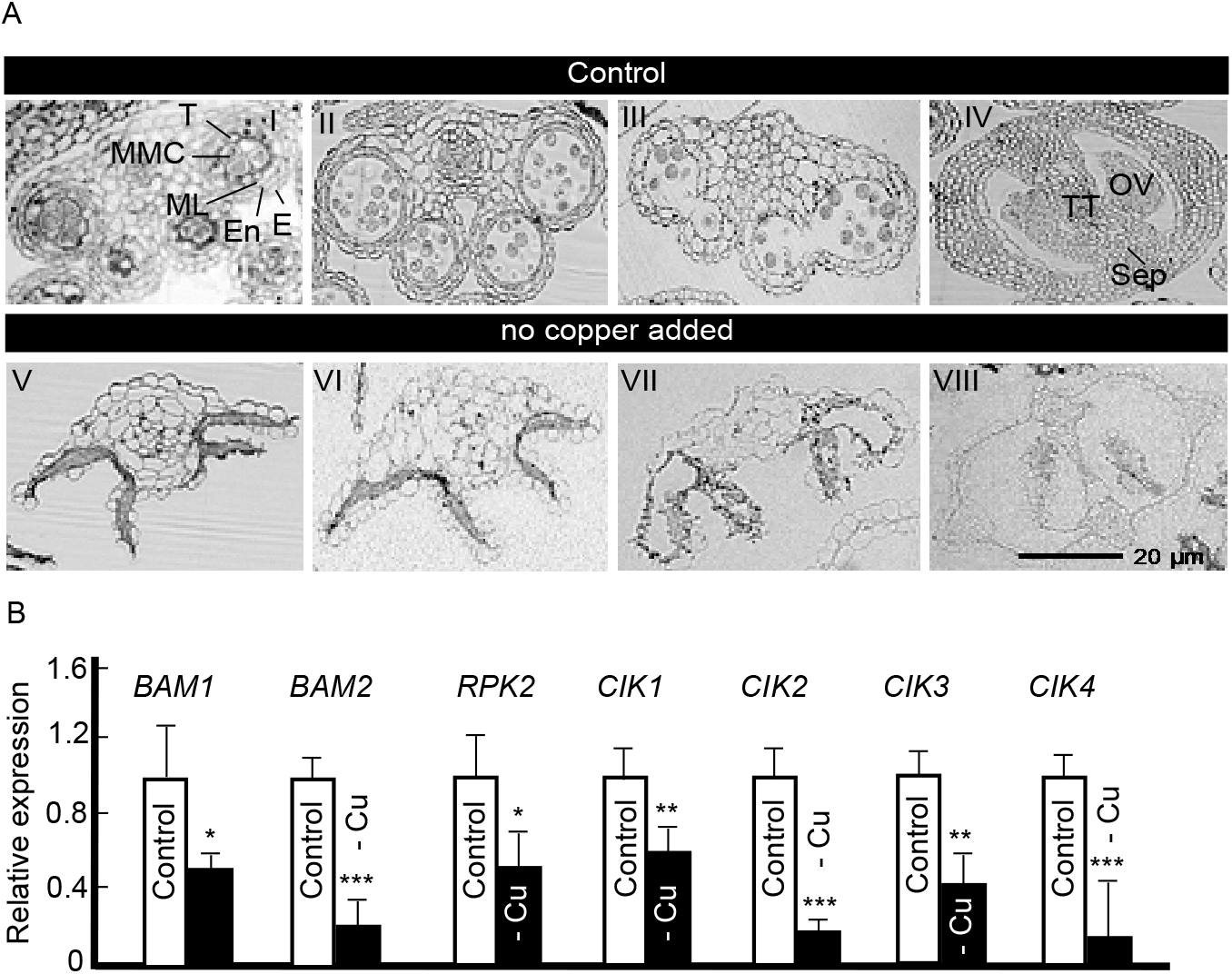
Copper deficiency causes malformation of both androecium and gynoecium during different developmental stages. (A) Representative images of the floral bud cross sections showing an anther and gynoecium. Plants were grown hydroponically with or without 250 nM CuSO_4_. I, II, III, IV are for plants grown in control conditions. V, VI, VII, and VIII are for copper-deficient plants. I and V are early stages of anther development, in which, microspore mother cells are present. II and VI are middle stages of anther development, in which, microspores are released. III and VII are late stages of anther development, in which, anthers are bilocular and contain tricellular pollen grains. IV and VIII are gynoecium cross sections present at late stages of anther development (i.e., III and VII). E, epidermis; En, endothecium; ML, middle layer; MMC, microspore mother cells; OV, ovule; Sep, Septum; T, tapetum; TT, transmitting tract. (B) Transcript abundances of anther specific genes in floral buds. Plants were grown hydroponically with or without 250 nM CuSO_4_. Values are mean ± SE (n = 3 independent experiments with at least 3-5 plants analyzed in each experiment). Asterisks, *, **, and *** indicate statistically significant differences compared to control condition with *P* < 0.05, 0.01, and 0.001, respectively, using the Student’s t-test.

### Copper deficiency decreases the expression of genes involved in early anther development

We then tested the effect of copper deficiency on the expression of genes associated with early stages of anther development. Specifically, we analyzed the expression of *BAM1* and *BAM2* (*Barely Any Meristem*), which encode CLAVATA1-related leu-rich repeat receptor-like kinases (LRR-RLKs) (Hord, Chen *et al*., 2006), *RPK2* which is also an LRR-RLK (Mizuno, Osakabe *et al*., 2007), and *CIK1* to *CIK4 (Clavata3 Insensitive Receptor Kinase1* to *4*) known as coreceptors of BAM1, BAM2, and RPK2 (Cui, Hu *et al*., 2018). RT-qPCR results showed that the transcript abundance of all marker genes was significantly reduced under copper deficiency compared to the control condition (**Fig. 5B**). These results were consistent with the observed anther developmental defects of copper-deficient plants (**Fig. 5A**) and provided another layer of evidence that copper deficiency causes malformation of reproductive organs.

### The expression of copper-microRNAs is upregulated in flowers of copper-deficient *Arabidopsis*

Transcript abundance of several microRNAs including *miR397, miR398, miR408*, and *miR857* is increased by a copper deficiency in roots and leaves of *A. thaliana* (Abdel-Ghany and Pilon, 2008; Pilon, 2017). Here, we tested whether the expression of these copper-microRNAs also increases under copper deficiency in *A. thaliana* flowers. To observe the dynamics and the specificity of their expression during flower development, we tested the expression of all copper-miRNAs in floral buds and open flowers. Among them, we detected the expression of *miR397a/b, miR398b/c* but not *miR398a*, and *miR857* (single gene) in floral buds but not in open flowers. On the other hand, we could detect *miR398a* and *miR408* (single gene) expression in open flowers but not in floral buds. RT-qPCR results showed that the transcript abundance of *c*opper-microRNAs tested, except for *miR408*, was significantly increased under copper deficiency (**Fig. 6A**), suggesting that they are involved in the response to copper deficiency in floral organs as well.

**Fig. 6.**
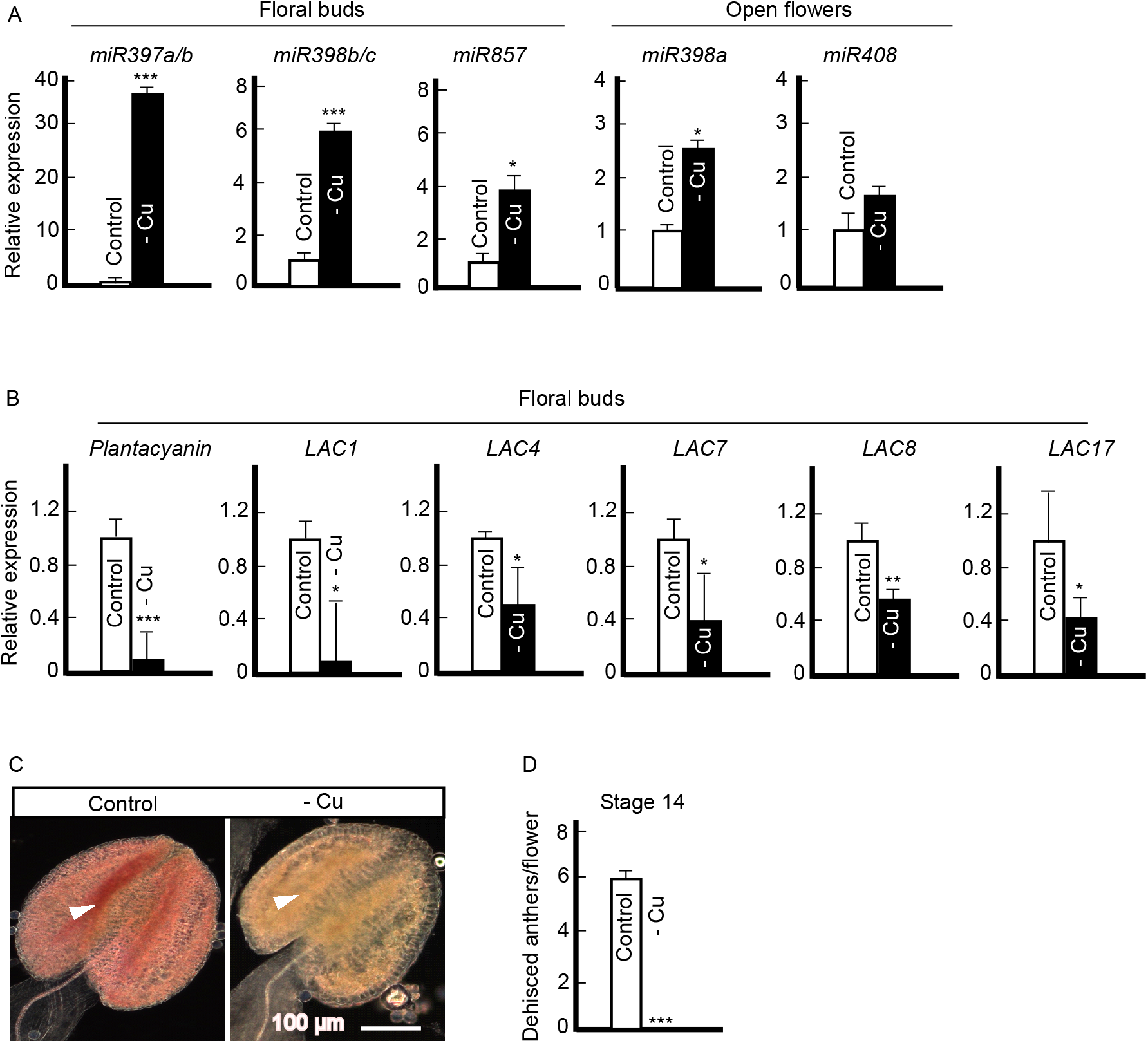
Copper deficiency upregulates the expression of copper miRNAs and reduces the expression of laccases and lignin accumulation. Transcript abundance of Cu-miRNAs (A), *plantacyanin* and lignin biosynthesis genes, *laccases*, (B) in open flowers or floral buds under copper deficiency. In (A) and (B), values are mean ± SE (n = 3 independent experiments with three to five plants analyzed in each experiment). Asterisks * (*P* < 0.05), ** (*P* < 0.001) and *** (*P* < 0.001) indicate statistically significant differences compared to control condition determined by the Student’s t-test. (C) A representative image of anther from plants grown with or without 250 nM copper. Phloroglucinol-HCl stains lignin in red. White arrowheads show stomium region in anther wall. Representative images are presented here from five independent experiments, each included at least 10 plants per condition. Note that copper-deficient anther lacks lignin in the stomium region in addition to everywhere else. (D) Dehisced anthers per single flower. Values are mean ± SE (n = 3 independent experiments). Thirty individual flowers from 10 individually grown plants without copper were analyzed. For control condition (250 nM CuSO_4_), 10 individual flowers from five individually grown plants were analyzed. Asterisks (***) indicate statistically significant difference compared to control condition with *P* < 0001 determined by the Student’s t-test.

### Copper deficiency reduces the expression of laccases in flowers of *Arabidopsis*

We next tested the effect of copper deficiency on the expression of some of the known copper-microRNA targets including *laccases (LAC1, LAC4, LAC7, LAC8*, and *LAC17*) encoding multicopper oxidases that are implicated in lignin synthesis and *plantacyanin*, encoding a plant-specific blue-copper protein (Dong *et al*., 2005; Nersissian *et al*., 1998; Zhang *et al*., 2018b; Zhao *et al*., 2013). We chose *LAC1, LAC4* and *LAC17*based on their decreased expression in our previous RNA-Seq data (Yan *et al*., 2017) in addition to *LAC7* and *LAC8* (Abdel-Ghany and Pilon, 2008). *miR397* targets *LAC4* and *LAC17*, while *miR408* and *miR857* target *plantacyanin* and *LAC7*, respectively. Although *LAC8* has no predicted *copper-microRNA* target sites, its expression is regulated by the copper supply (Abdel-Ghany and Pilon, 2008). We found that the transcript abundance of all *LAC* genes and *plantacyanin* was significantly decreased under copper deficiency in floral buds (**Fig. 6B**). We also tested the effect of copper deficiency on *Cu-microRNAs* and their corresponding targets in the primary inflorescence. The transcript abundance of all tested microRNAs and all tested *LAC* genes was significantly up- and down-regulated, respectively, by a copper deficiency in primary inflorescences (**Supplementary Fig. S2**).

### Copper deficiency reduces lignin accumulation in anthers and reduces anther dehiscence

Because the transcript abundance of several *LAC* genes including *LAC4* and *LAC17* was reduced in flowers under copper deficiency, we predicted that lignin deposition will be decreased in copper-deficient plants as well. We found a dramatic reduction of lignin staining in anthers and primary inflorescence of copper-deficient plants (**Fig. 6C** and **Supplementary Fig. S2C**). Lignin deposition was observed in the xylem, including the vessels, parenchyma, and the interfascicular region of inflorescence stems of plants grown under copper sufficiency. However, lignification was completely abolished in the interfascicular fibers and was only detectable in the xylem vessels in copper-deficient plants (**Supplementary Fig. S2C**). Concerning anthers, lignin staining was observed in the stomium regions of plants grown under copper replete conditions while was nearly absent in these regions in plants grown under copper deficiency (**Fig. 6C**, white arrowhead). Consistent with the important role of lignification and associated anther wall thickening in anther dehiscence (Mitsuda *et al*., 2005), nearly 100% of anthers from copper-deficient plants were indehiscent (**Fig. 6D**).

### The expression of senescence-associated genes, *SAG12, SAG13*, and *WRKY53* is altered in young and mature leaves in copper-deficient *Arabidopsis*

Our recent studies in *A. thaliana* have shown that copper deficiency triggers the foliar accumulation of jasmonic acid (JA) (Yan *et al*., 2017). Because JA, among its other physiological functions, is also considered as one of the early signals stimulating leaf senescence, we hypothesized that copper deficiency may trigger leaf senescence as well. To test our hypothesis, we evaluated the effect of copper deficiency on the expression of the senescence-associated genes that are also the downstream of JA targets, *SAG12, SAG13*, and *WRKY53* (Woo *et al*., 2019). Because young leaves are more susceptible to copper deficiency than mature leaves due to poor copper phloem-based mobility (Broadley *et al*., 2012), we anticipated that young leaves might display more dramatic molecular responses of senescence. Consistent with our hypothesis, we found that the expression of *SAG12* was upregulated by nearly 2- and 17-fold in mature and young leaves, respectively under copper deficiency (**Fig. 7**). Surprisingly, we found that the expression of *SAG13* and *WRKY53* was significantly downregulated in mature leaves of copper-deficient *vs*. copper-sufficient plants (**Fig. 7**). The expression of *WRKY53* was also significantly downregulated in young leaves of copper deficient *vs*. copper-sufficient plants (**Fig. 7**). These data show that copper deficiency mounts a distinct transcriptional response of senescence-associated genes, and perhaps, triggers distinct aspects of senescence.

**Fig. 7.**
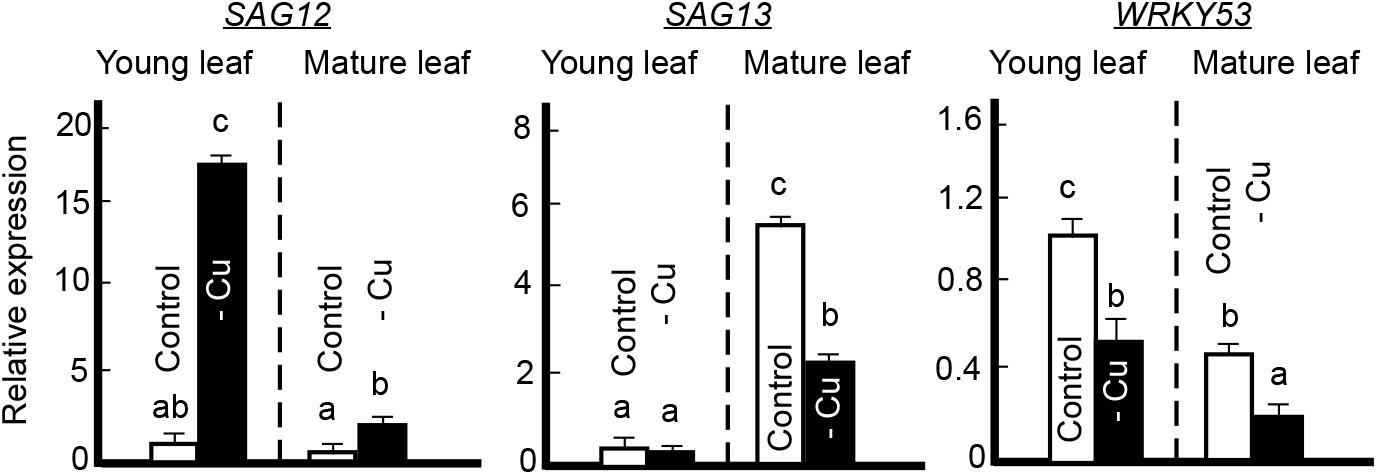
Copper deficiency decreases the expression of *SAG13* and *WRKY53* but triggers the expression of *SAG12* upon transition to bolting. Expression of *SAG12, SAG13*, and WRKY53, as senescence marker genes, in young and fully mature green rosette leaves upon bolting. Plants were grown hydroponically with or without 250 nM CuSO_4_ until bolting. The approximate plants age were five to six weeks. Shown are mean values ± SD of a representative experiment of three independent experiments. Each experiment analyzed three to five plants. Lowercase letters are used to indicate statistically significant differences between conditions (*P* < 0.05) determined by the Tukey-Kramer HSD test using JMPPro 14. Data were normalized to control in a young leaf.

## Discussion

The important role of the micronutrient copper in plant fertility has been recognized for more than 30 decades. A range of crop species, including wheat, oat, barley, sweetcorn, and sunflower was used to show that copper deficiency affects reproduction more strongly than vegetative growth, may delay flowering and leads to male sterility (Dell, 1981; Graham, 1978; Graves and Sutcliffe, 1974). These pioneering studies have been very informative but yet scattered and raised more mechanistic questions including what are the sites of copper accumulation in flowers, which aspects of copper metabolic functions are important for ensuring successful fertility, which transport pathways are responsible for copper delivery to reproductive organs and how these transport pathways are regulated. In addition, a systematic and comprehensive analysis of the effect of copper deficiency on plant fitness with a focus on its impact on fertility has not been yet conducted. Here, we used a model dicot *A. thaliana* to initiate a comprehensive analysis of molecular and mechanistic reasons underlying the copper-deficiency mediated late flowering, poor pollen germination and compromised reproduction.

### Copper deficiency alters shoot architecture in Arabidopsis

While establishing the range of copper concentrations that can be regarded as adequate for growth of *A. thaliana* in hydroponics, we noted that plants grown without or with low (5 nM) copper supplementation had small rosette size (**Supplementary Fig. S1**). This finding was expected because of the essential role of copper in respiration and photosynthesis. In fact, about 50% of copper found in plants is present in chloroplasts, where it is bound to plastocyanin, a copper-containing protein that mediates electron transfer between PSII and PSI (Weigel *et al*., 2003). Therefore, copper-deficient plants have low rates of photosynthesis and reduced carbohydrate production that, in turn, is reflected in the reduced plant growth and development (Broadley *et al*., 2012; Brown and Clark, 1977; Ravet and Pilon, 2013a). In addition, the reduced respiration-based energy supply for the energy-dependent processes are among the contributing factors for the decline of growth rates and failure to reach the full-size potential of the developing leaf. With that notion, it was surprising to find that copper-deficient plants had longer primary inflorescence upon transition to flowering and developed more axillary branches (**Fig. 1A, B**).

It is recognized that the plasticity of shoot architecture depends on differential activation of axillary buds, environmental conditions and interactions between systemically moving phytohormones auxin, strigolactones, and cytokinins (reviewed in (Teichmann and Muhr, 2015), (Wang *et al*., 2018) and (Domagalska and Leyser, 2011)). Among these phytohormones, the prominent role of auxin in exerting apical dominance is well established. Removal of the shoot apex (decapitation) results in lateral bud activation and shoot branching, while reapplication of exogenous auxin to the stump restores branching inhibition (reviewed in (Domagalska and Leyser, 2011). In our experiments, we noted that copper deficiency leads to the abortion of apical flower buds (**Fig. 1C**). Thus, we speculated that copper deficiency promotes shoot branching via mitigating the apical dominance effect and altering auxin signaling. Consistent with this suggestion, the reapplication of IAA or copper to the shoot apex partially rescued the branching phenotype of copper-deficient plants (**Fig. 1C**). It is noteworthy that copper but not auxin reapplication to the shoot apex partially restored seed (**Fig. 1C**). The relationship between copper and auxin homeostasis remains to be elucidated but it is noteworthy that copper promotes auxin accumulation and cell proliferation in the copper moss *Scopelophila cataracta* (Nomura *et al*., 2015), and excess copper prevents auxin redistribution in the root through interacting with an auxin efflux carrier, Pinformed1 (PIN1), (Yuan *et al*., 2013). The precise nature of documented here copper-deficiency promoted changes in shoot architecture remains to be established.

### Copper deficiency delays vegetative to reproductive stage transition by reducing the expression of *FT*

To ensure successful reproduction, plants control their flowering time by changing their growth rates and/or altering the vegetative-to-reproductive stage transition (Cho *et al*., 2017; Schmalenbach *et al*., 2014; Simpson and Dean, 2002). Although it is accepted that poor nutrition tends to promote flowering, low phosphorus and nitrogen have distinct effects on flowering time in *A. thaliana* (Cho *et al*., 2017). Specifically, nitrate□limiting conditions promote flowering independently of light, gibberellin and autonomous pathways (Castro Marín *et al*., 2011). By contrast, phosphorus deficiency delays flowering (Kant *et al*., 2011). Our observations presented here and in Sheng et al, 2019 show that copper deficiency delays flowering in *A. thaliana* and *B. distachyon*. It is unclear, however, whether delayed flowering under copper deficiency is a result of slower growth rates due to reduced photosynthesis and respiration and/or a delayed developmental transition from the vegetative-to-reproductive stage.

We found that copper deficiency delayed flowering and increased accumulation of rosette leaves in *A. thaliana* (**Fig. 2A, B**). This finding is consistent with the conclusion that copper deficiency does not simply reduce the plant growth rates but impacts the developmental transition from the vegetative-to-reproductive stage. While the specific role of copper in developmental transition is yet to be established, it is possible that plants stay longer in the vegetative stage to accumulate the critical level of photosynthates. In line with this suggestion are past studies showing that photosynthetic activity influences flowering (Bernier *et al*., 1993) and that *A. thaliana* exposed to strong irradiation flowers sooner, and has the increased levels of endogenous sucrose in leaves (King *et al*., 2008). Sucrose also promoted flowering in several species, and the exogenous application of a low concentration of sucrose partially rescues the late□flowering phenotypes of *Arabidopsis* mutants (Bernier *et al*., 1993; Cho *et al*., 2018; Ohto *et al*., 2001). Trehalose-6-phosphate has been implicated in the regulation of flowering time in *A. thaliana* and the downregulation of trehalose-6-phosphate synthase expression significantly delayed flowering even though the basal sucrose level remains unchanged (Cho *et al*., 2018; Wahl *et al*., 2013). It is suggested that sucrose functions in the leaf phloem while trehalose-6-phosphate functions in the shoot apical meristem to enhance the generation of florigens such as *FT* (Cho *et al*., 2018; Wahl *et al*., 2013). Specifically, the increased endogenous sucrose levels due to higher photosynthetic activity lead to higher expression of *FT*, hence, sucrose-mediated signals are regarded to function upstream of *FT* and are intimately related to the plant photosynthetic capacity (King *et al*., 2008; Seo *et al*., 2011). In this regard, it is noteworthy that the transcript abundance of *FT* was significantly decreased in leaves of *A. thaliana* under copper deficiency (**Fig. 2D**). Thus, it is possible that copper deficiency delays flowering time indirectly *via* reducing photosynthetic rates, decreasing the sucrose and perhaps, trehalose-6-phosphate level, which in turn, leads to the reduction of *FT* expression (**Fig. 8**).

**Fig. 8.**
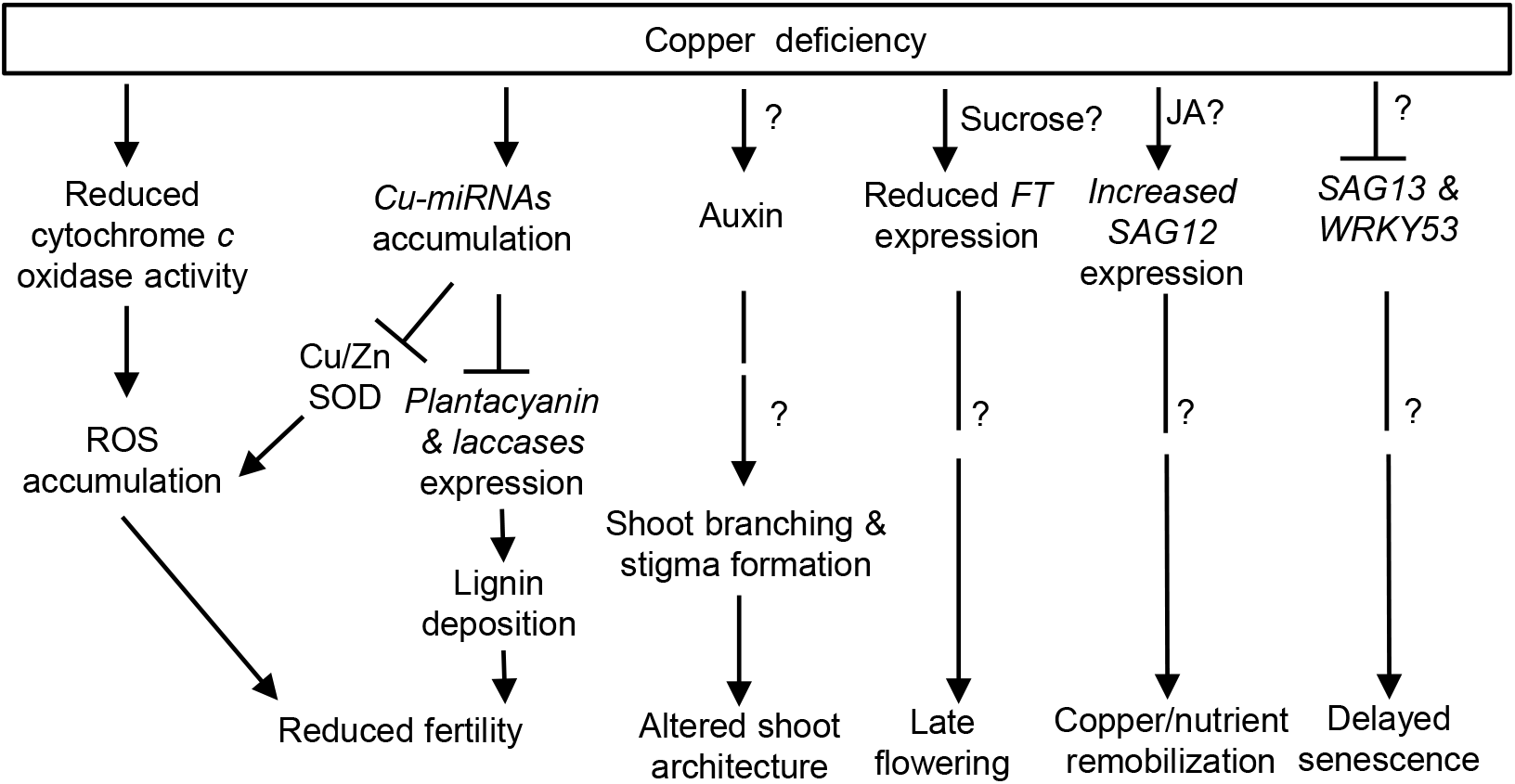
Proposed model of the effect of copper deficiency on development and reproduction of *Arabidopsis thaliana*. Under copper deficiency, *copper miRNAs* (*Cu-miRNAs*) accumulate leading to a degradation of their targets, i.e., plantacyanin and laccases. The reduced level of plantacyanin and laccases, contributes to the reduced fertility, due to the failure in anther dehiscence and pollen-stigma interaction. Copper deficiency changes shoot branching and stigma formation possibly via changes in the auxin level, leads to altered shoot architecture and reduced fertility, respectively. Further we showed copper deficiency resulted in accumulation of ROS in pollen, leading to imbalance in cellular redox status, which in turn, causing reduced fertility. This accumulation of ROS could be the result of reduced enzyme activity of either cytochrome *c* oxidase or Cu/Zn-SOD or both under copper deficiency. It is noteworthy that the reduction in the cytochrome *c* oxidase enzyme activity also decreases the ATP level, which in turn, could lead to the reduced pollen germination, and thus reduced fertility. Further, copper deficiency resulted in the reduced expression level of *FT* gene, causing late flowering. Sucrose is one possible contributor acting upstream of *FT*. Furthermore, we showed copper deficiency triggers accumulation of *SAG12* transcript, possibly through jasmonic acid (JA), suggesting its involvement in the copper/nutrient remobilization. On the other hand, reduced expression of *SAG13* and *WRKY53* under copper deficiency suggests overall delay in developmental senescence. Question marks (?) indicate components of the pathway not yet experimentally characterized.

### Copper deficiency increases the expression of *miR172* in leaves of *A. thaliana*

Among factors mediating the transition to flowering in *A. thaliana* is a conserved regulatory module including two microRNA families, miR156 and miR172, and their corresponding downstream targets. The transition to flowering is associated with the reduction of *miR156* accumulation and a concomitant increase in *miR172* and *FT* (Teotia and Tang, 2015; Wang *et al*., 2009). Because we found that the transcript abundance of *FT* is reduced in leaves of copper-deficient plants (**Fig. 2D**), we anticipated that the expression of *miR172* will be reduced as well. Unexpectedly, we found that the transcript abundance of both *miR172c* and *d* isoforms that we were able to detect in leaves of *A. thaliana*, increased significantly under copper deficiency (**Fig. 2E**). Why the upregulation of *miR172* did not lead to the upregulation of *FT* and concomitant transition to flowering, is not clear. It is possible that the miR172-stimulated *FT* expression is overridden by severe copper deficiency that, in turn, is expected to reduce the photosynthetic capacity, sucrose accumulation and *FT* expression (**Fig. 8**). It is also possible that copper deficiency-mediated increase in the transcript abundance of *miR172* is not sufficient to promote floral induction under copper deficiency or that copper deficiency may also activate floral repressors including *Flowering Locus C* (*FLC*) (Crevillen and Dean, 2011; Ortuño-Miquel *et al*., 2019), which in turn, will inhibit *FT* expression and delay flowering, independently of *miR172*. Future studies will establish the interactions between copper deficiency and pathways controlling flowering time.

### Copper deficiency reduces fertility and causes defects in both reproductive organs

Copper deficiency-mediated defects in reproduction have been linked to male infertility (Dell, 1981; Graham, 1978). Our recent x-ray synchrotron-fluorescence microscopy analysis of the spatial distribution of copper in flowers of *A. thaliana* identified bulk of copper in anthers (Yan *et al*., 2017). Failure to deliver copper to anthers in the mutant lacking two regulators of copper homeostasis, SPL7 and CITF1, resulted in a dramatic loss of fertility (Yan *et al*., 2017). However, Yan et al., 2017 have also shown that in addition to anthers, copper is associated with pistils, suggesting that this mineral can function in female fertility as well. Consistent with this suggestion results from reciprocally crossed *A. thaliana* with one parent grown under copper sufficiency while another under copper deficiency, implicated copper in the fitness of both male and female gametophytes (**Table 1**). Moreover, we showed that the gynoecium is much more impacted by copper deficiency than androecium. This was evidenced by the failure of plants to produce seeds when the female of a copper-deficient plant was a recipient of pollen from a copper-sufficient plant. By contrast, plants grown under copper sufficiency and fertilized with pollen from the copper-deficient plant still produced some seeds, albeit at a significantly reduced level (**Table 1**).

Further, we noticed that stigmas of *A. thaliana* grown under copper deficiency had either reduced papillae formation or had severely reduced papillae length (**Fig. 4A**). Because metabolically active papillar cells are required for successful pollen tube growth (Kandasamy *et al*., 1993), the infertility of gynoecium of copper-deficient plants might be, in part, caused by a defect in papillae formation. It is noteworthy that the defect in copper delivery to reproductive organs in the mutant lacking copper transporter YSL3 also leads to aberrant stigma formation in *B. distachyon (Sheng et al., 2019)*. Why copper deficiency causes defects in stigma morphology is unclear. Mounting evidence suggests that auxin plays an important role in apical–basal patterning that includes stigma and style patterning of the gynoecium. For example, the reduced development of the stigmatic papillae observed in *A. thaliana* mutants for the *bHLH* family member *SPATULA* (*SPT*) can be rescued by blocking auxin transport using naphthylphthalamic acid (NPA) or by inducing the expression of the *STYLISH1* (*STY1*) that activates the auxin biosynthesis gene *YUCCA4* (Nemhauser *et al*., 2000; Sohlberg *et al*., 2006) and reviewed in (Marsch-Martínez and de Folter, 2016). Since our findings link copper deficiency-promoted shoot branching defects to auxin (**Fig. 1**), it is tempting to speculate that copper deficiency-caused defect in papillae formation is linked to auxin homeostasis as well.

Results presented here also substantiate the role of copper in male fertility. Specifically, we show that copper deficiency significantly reduced the number of pollen grains, pollen viability, and germination in *A. thaliana* (**Fig. 4B, D**). These results are consistent with past findings showing that anther and pollen development are sensitive to copper status (Agarwala, Sharma *et al*., 1980, Jewell, Murray *et al*., 1988, Pandey 2010). Graham (1975) reported that male sterility of wheat occurs near the meiosis stage of pollen mother cells. Using crosspollination, this group has shown that the copper deficiency-caused infertility is due to pollen sterility but not due to the female gametophyte. This contrasts with our findings in *A. thaliana*, in which, cross-pollination results showed that female fertility is entirely reduced under copper deficiency (**Table 1**). This discrepancy may be due to the species-specific sensitivity to copper deficiency.

The defect in pollen fertility was also observed in *A. thaliana, B. distachyon*, and *Oryza sativa* mutants lacking transporters that mediate copper delivery to reproductive organs (Chu *et al*., 2010; Sheng *et al*., 2019; Zhang *et al*., 2018a). We show that adding copper directly to the pollen germination medium rescued pollen germination defect of copper-deficient *A. thaliana* (**Fig. 4F**). This is consistent with the previous study in rice, where copper application rescued pollen germination defect of the rice mutant lacking copper-nicotianamine transporter, OsYSL16 (Zhang, Lu *et al*., 2018). Here we also show that copper deficiency-caused defect in pollen germination is, in part, caused by the altered redox status of the pollen grains and could be linked to the reduced cellular energy levels. First, we showed that pollen grains from copper-deficient plants accumulated a high level of ROS while supplementing pollen germination medium with an antioxidant, L-ascorbate, rescued pollen germination defect of copper-deficient plants (**Fig. 4E, F**). Second, COX activity was significantly reduced under copper deficiency (**Table 2**). The reduction in COX activity results in over-accumulation of O_2_ in the mitochondria matrix, which, in turn, leads to over-production of ROS and consequent damage to cells. ROS accumulation under copper deficiency can also stem from the reduced Cu/Zn SOD activity (Ravet and Pilon, 2013b). In addition, the reduction in COX activity leads to a reduction in cellular ATP production (Droppa *et al*., 1984). Consistently, *A. thaliana* COX11 homolog is involved in the insertion of copper into the COX complex during its assembly in mitochondria, is expressed in germinating pollen among other tissues, and its loss-of-function impairs pollen germination (Radin *et al*., 2015). Together, our results implicate an imbalance in the ROS level and the reduced energy level in copper-deficiency mediated male infertility.

### Copper deficiency leads to abnormal anther development

We further detected anther abnormality in copper-deficient plants (**Fig. 5A**). Anthers are the sites of pollen development. At the early stage of anther development (anther stage 5, floral stage 9), four cell layers are present in the anther wall including epidermis, endothecium, middle layer, and tapetum that is a nutritive tissue (**Fig. 5A-I)**. At this stage, pollen mother cells (PMCs) are formed within anther locules. PMCs undergo meiosis to form microspores, which are then differentiated into the three-cell pollen grains at stage 12 of anther and flower development (Sanders *et al*., 1999). Under copper deficiency, however, except for epidermis, the other three cell layers, including tapetum, were absent from the anther wall (**Fig. 5A-V**). These undefined structures remained throughout the middle and late stages of anther development in copper-deficient anthers (**Fig. 5A-VI and -VII**). Similarly, gynoecium cross-sections of the copper-deficient flowers also showed undefined structures (**Fig. 5A-VIII**). These data suggest that copper deficiency leads to the defects at early stages of anther development.

We speculate that these defects in copper-deficient flowers occur due to the significantly reduced expression of genes involved in anther cell determination (**Fig. 5B**). Specifically, Arabidopsis, *BAM1* and *BAM2* (*Barely Any Meristem*) encode CLAVATA1-related leu-rich repeat receptor-like kinases (LRR-RLKs). The loss-of-function of both genes in the *bam1bam2* double mutant shows abnormal anther lacking the endothecium, middle, and tapetum layers (Hord, Chen *et al*., 2006). *RPK2* is also an LRR-RLK which is required for early anther development (Mizuno, Osakabe *et al*., 2007). Anthers in the *rpk2* mutant lack the middle layer, have abnormal hypertrophic tapetal cells, as well as thickened and lignified endothecium cells, which together, lead to failure in pollen production and release. CIK1 to CIK4 (Clavata3 insensitive receptor kinase1 to 4) have shown to function as coreceptors of BAM1 and BAM2 and RPK2 to control cell fate specification during early anther development in Arabidopsis (Cui, Hu *et al*., 2018). The loss-of-function of these four CIKs reduces fertility.

### Copper deficiency-based reduction in anther dehiscence is mediated, in part *via* the reduced lignification, which in turn, may occur through conserved copper-miRNAs

The expression of copper microRNAs including *m/R397, miR398, miR408*, and *miR857*, is upregulated in roots and leaves of *A. thaliana* in response to copper deficiency to facilitate the turnover of non-essential copper proteins and contribute to the copper economy (Pilon, 2017). Here, we show that copper microRNAs respond to copper deficiency differently depending on the stage of flower development. Specifically, *miR397a/b, miR398b/c*, and *miR857* were upregulated in floral buds but not in open flowers, while *miR398a* and *miR408* were upregulated in open flowers but not in floral buds. This observation suggests the unique roles of specific copper-microRNAs in maintaining copper homeostasis during flower development.

Among well-established copper microRNAs targets are copper proteins, *plantacyanin* and *laccases* (Pilon, 2017). Thus, it was not surprising that the expression of *plantacyanin* and all *LAC* genes tested was downregulated in flowers and primary inflorescence (**Fig. 6; Supplementary Fig. S2**). Plantacyanins have a conserved copper-binding site and are not essential for the growth and development of plants as evidenced by a lack of defects in *A. thaliana* and *O. sativa* plantacyanin mutants (Dong *et al*., 2005; Einsle *et al*., 2000; Ryden and Hunt, 1993; Zhang *et al*., 2018b). Thus, the decrease in *plantacyanin* expression under copper deficiency is part of the copper economy mode aiming to channel copper from non-essential to essential copper proteins.

Consistent with the role of *LAC4, LAC11*, and *LAC17* in lignin deposition in *A. thaliana* roots and stems (Zhao *et al*., 2013), the downregulation of their expression under copper deficiency (**Fig. 6B**) was associated with the decreased lignin accumulation in anthers and primary inflorescence (**Fig. 6C; Supplementary Fig. S2**). Secondary wall thickening that includes lignification and cellulose deposition, occurs in the endothecium layer in anthers and is essential for anther dehiscence and pollen dispersal (Cecchetti *et al*., 2013; Hao *et al*., 2014a; Mitsuda *et al*., 2005; Yang *et al*., 2017). Thus, we propose that the entirely abolished anther dehiscence in copper-deficient plants (**Fig. 6D**) is associated with the loss of lignin in the endothecium and stomium region (**Fig. 6C**) and is among contributing factors of the reduced fertility under copper deficiency. Lignin deposition was completely abolished from the interfascicular fibers and was only associated with the xylem (**Supplementary Fig. S2**). While interfascicular fibers are not essential for plant survival, their development and lignification are important for the stem strength and structural support of the plant body (Zhong *et al*., 1997). Failure to lignify this tissue results in weakening of stems that ultimately can translate to crop lodging in the field (Broadley *et al*., 2012; Zhong *et al*., 1997).

### The relationship between copper deficiency and leaf senescence

Natural leaf senescence is an important developmental process that ensures the remobilization of nutrients including minerals from the senescing tissues to developing reproductive organs and seeds (Leopold, 1961). Natural leaf senescence which is typically triggered by the age of leaves, is associated with a transition to reproduction and can be initiated by hormones including jasmonic acid (JA)(Woo *et al*., 2019). Environmental stresses, as well as nutrient deficiency, are known to cause premature senescence (Leopold, 1961; Woo *et al*., 2019; Xie *et al*., 2014). While natural senescence increases reproductive success, premature senescence is often correlated with severely decreased yields (Leopold, 1961; Woo *et al*., 2019; Xie *et al*., 2014). We have shown recently JA levels increase in leaves of copper-deficient *A. thaliana* suggesting that deficiency for this mineral causes premature senescence, and could be among the reasons for dropped seed yield (Yan *et al*., 2017). On the other hand, we and others have shown that copper deficiency delays the transition to flowering (**Fig. 2**) and thus, would be expected to delay senescence. Hill et al., 1978 have also shown that copper deficiency delays chlorophyll degradation of mature wheat leaves, further reinforcing the notion that unlike other mineral deficiencies that trigger senescence, copper deficiency might delay it.

To reconcile this discrepancy, we evaluated the effect of copper deficiency on the expression of senescence marker genes, *SAC12, SAG13*, and *WRKY53. SAG12* encodes a cysteine protease, which is expressed only in senescing tissues. It is involved in nitrogen remobilization and its expression is upregulated by JA (He *et al*., 2002; James *et al*., 2018; Noh and Amasino, 1999). *SAG13* encodes a senescence-associated protein that is also required for resistance against fungal pathogens. *SAG13* is induced during stresses such as oxidative stress and is required for the normal seed germination, seedling growth, and anthocyanin accumulation (Dhar *et al*., 2020). *WRKY53* is a master regulator of age-induced leaf senescence that is associated with the onset of natural senescence and acts upstream in the senescence transcriptional cascade (Zentgraf *et al*., 2010). Both *SAG12* and *SAG13* are downstream WRKY53 targets with *SAG12* considered a marker for age-dependent natural senescence while *SAG13* is commonly used to evaluate stress-induced senescence (Zhao *et al*., 2018).

We found that copper deficiency mounted a distinct transcriptional response of senescence-associated genes (**Fig. 7**). Specifically, while the expression of *SAG12* was significantly upregulated by a copper deficiency in both mature and young leaves, the expression of *SAG13* and *WRKY53* was significantly downregulated in leaves of copper-deficient plants (**Fig. 7**). Given that WKRY53 acts upstream in the transcriptional cascade leading to leaf senescence and its expression is downregulated under copper deficiency, we conclude that unlike other stress factors, copper deficiency delays the onset of senescence. This suggestion is also consistent with the delayed transition to flowering under copper deficiency (**Fig. 2**). However, the question remains why *SAG12* that is also among WRKY53 targets is upregulated under copper deficiency and why its transcript level is nearly 9-fold higher in young *vs*. mature leaves in copper-deficient *vs*. copper-sufficient plants. The increased expression of *SAG12* under copper deficiency may be triggered by JA, which, accumulates in leaves of copper-deficient plants (He *et al*., 2002; Yan *et al*., 2017). It is also possible that a nutritionally-derived, yet unidentified, signal triggers the upregulation of *SAG12* expression independently of WRKY53 to initiate nutrient remobilization from leaves to, eventually, ensure the transition to flowering. In this regard, copper deficiency is more pronounced in young leaves because of poor phloem-based copper mobility (Broadley *et al*., 2012). Coincidentally, the *SAG12* expression is highly upregulated by the copper deficiency in young leaves, reinforcing the notion that nutritionally derived signal may exist to trigger the *SAG12* expression for stimulating protein degradation and amino acid utilization required for the growth of young leaves and subsequent transition to flowering.

To conclude, copper deficiency exerts pleiotropic effects on plant growth and development that include auxin-related changes in shoot architecture, delayed transition to flowering through the decreased expression of *FT*, possibly *via* reducing the production of carbohydrates including the sucrose, the established *FT* activator (**Fig. 8**). Copper deficiency also delayed senescence as evidenced by the significantly decreased expression of the master regulator of the onset of senescence, *WRKY53*. Copper deficiency triggered *SAG12* expression *via* the WRKY53-independent pathway that, we hypothesize, involves copper deficiency-derived nutrient signaling pathway. The reduced pollen fertility of copper-deficient plants stems from multiple factors including the failure of anthers to dehisce and disperse pollen as well as the increased ROS level concomitant with the decreased cellular energy (**Fig. 8**). Copper-deficiency gynoecium infertility can be, in part, caused by reduced stigma papillae formation. Overall, this study opens several new areas for the in-depth investigation into the relationship between copper homeostasis and hormone-mediated shoot architecture, gynoecium fertility and copper deficiency-derived signals leading to the delay in flowering time and senescence.

## Supplementary data

Supplementary data are available at JXB online.

---

**Supplementary Table S1.**
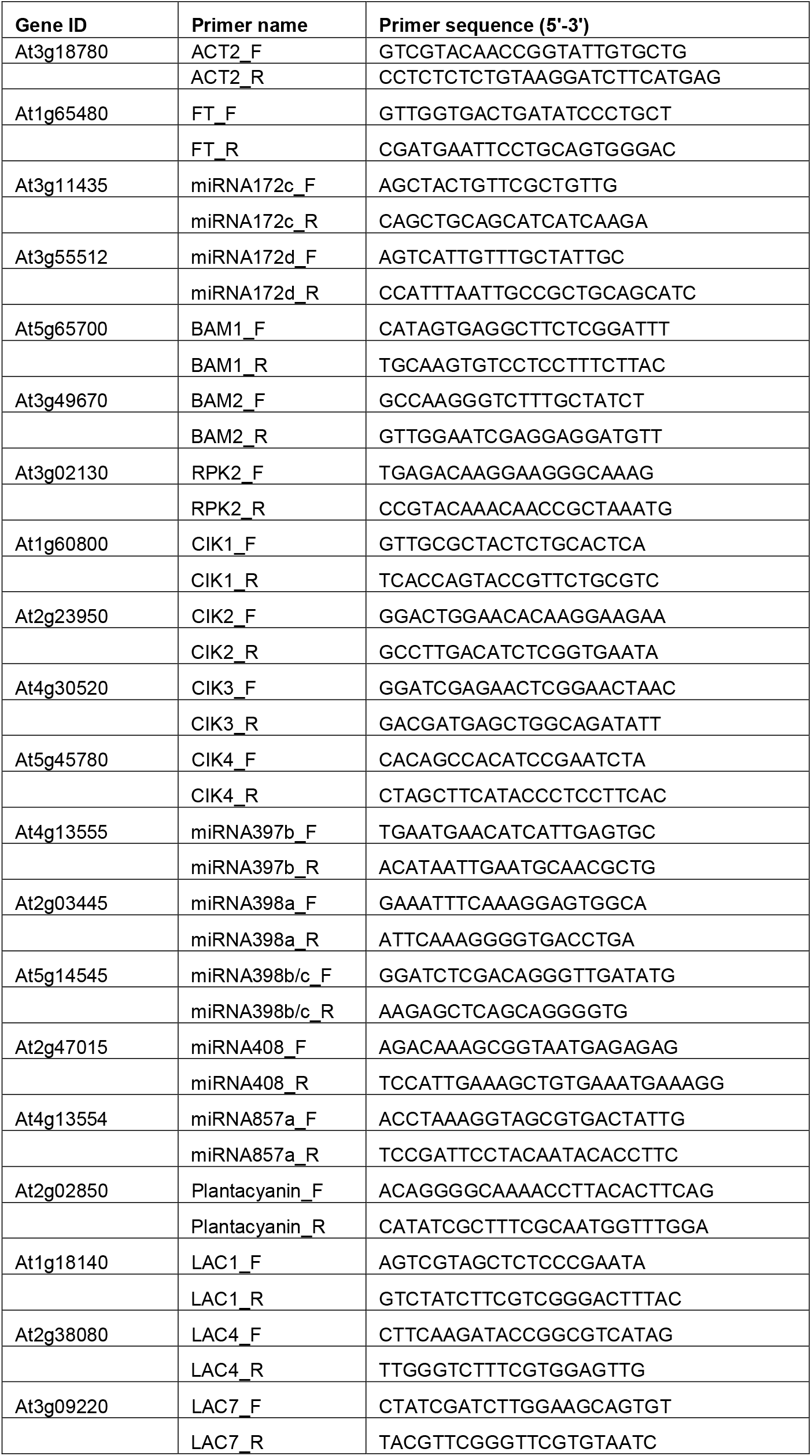
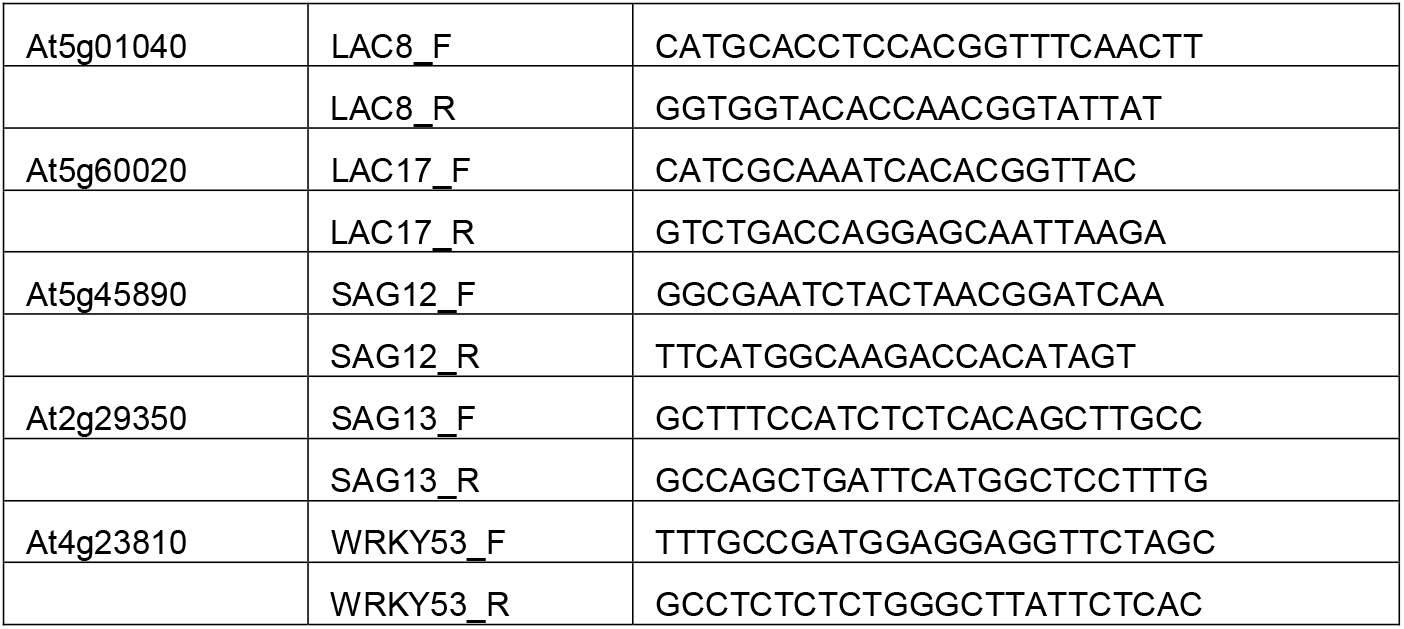
List of RT-qPCR primers and their sequences.

**Supplementary Fig. S1. Copper deficiency leads to small stature in Arabidopsis.**

(A) Shows representative images of plants grown hydroponically with or without 250 nM CuSO_4_ at flowering. Scale bar = 1 cm. (B) Rosette leaf length is shown upon transition to flowering for plants grown hydroponically under different copper concentrations, as indicated. Values are mean ± SE (n = 3 independent experiments with 5-10 plants analyzed in each experiment). *** indicates statistical significance difference compared to control condition with *P* < 0.0001 using the Student’s t-test.

**Supplemental Fig. S2. Copper deficiency decreases lignin biosynthesis in primary inflorescence through accumulation of copper miRNAs.**

(A) Increase in the transcript abundances of copper miRNAs in inflorescence stem under copper deficiency. (B) Decrease in the transcript abundances of lignin biosynthesis genes, laccases, under copper deficiency. (A) to (B) Plants were grown hydroponically with or without 250 nM CuSO_4_ until flowering. * (*P* < 0.05), ** (*P* < 0.001) and *** (*P* < 0.001) indicate statistically significant differences compared to control condition determined by the Student’s *t*-test. Values are mean ± SE (n = 3 independent experiments with at least three to five plants analyzed in each experiment). (C) Cross section of primary inflorescence showing lignin deposition pattern. Phloroglucinol-HCl stains lignin in red. Representative images are presented here from three independent experiments, each with at least 10 plants per condition. P and X refer to phloem and xylem, respectively. White arrowheads show site of lignin deposition in the interfascicular xylem fibers which is abolished under copper deficiency condition.

## Acknowledgements

We thank John Grazul and Josh Strable at the Cornell Center for Materials Research (CCMR) and Michael Scanlon lab, respectively, for assistance in preparing Ultra-thin sections from flower buds. This study was funded by NSF-IOS #1656321 awarded to O.K.V.

## Abbreviations

BAM1: Barely any meristem 1
BAM2: Barely any meristem 2
CIK1 to 4: Clavata3 insensitive receptor kinase1 to 4
CITF1: Copper deficiency induced transcription factor 1
Cu: Copper
Cu-miRNA: Copper microRNA
Cu/Zn-SOD: Copper/zinc-superoxide dismutase
COX: Cytochrome *c* oxidase
FT: Flowering locus T
JA: Jasmonic acid
LAC: Laccase
miRNA: MicroRNA
ROS: Reactive oxygen species
RT-qPCR: Real time quantitative polymerase chain reaction
SAG12: Senescence-associated gene 12
SAG13: Senescence-associated gene 13
SPL7: Squamosa promoter binding protein–like7
WT: Wild type
ZT: Zeitgeber time

